# Optimizing survey methods for spiders and harvestmen assemblages in an Amazonian upland forest

**DOI:** 10.1101/093740

**Authors:** Ana Lúcia Tourinho, Sidclay C. Dias, Nancy F. Lo-Man-Hung, Ricardo Pinto-da-Rocha, Alexandre B. Bonaldo, Fabricio B. Baccaro

## Abstract

Invertebrates can be sampled using any of several well-established, rapid and cost-effective methods for documenting species richness and composition. Despite their many differences, different orders of arachnids have been often sampled together in various studies. Active nocturnal search has been long considered the most efficient method for sampling spiders and harvestmen in tropical forests. We compared the number of species and composition of spiders and harvestmen simultaneously sampled using three sampling methods: beating tray, active nocturnal search and Winkler traps at areas along the Urucu River, Coari, Amazonas. We found that a reasonable inventorying of harvestmen can be accomplished solely by nocturnal search, whereas the beating tray and Winkler approaches are redundant. For spiders, both the nocturnal and beating tray methods were complementary and are needed to provide a more complete picture of spider assemblages. An inventory based solely on nocturnal search saves 75% of the survey costs for harvestmen assemblages and 46% for spider assemblages. Based on our findings we propose that different taxonomic groups (e.g. harvestmen and spiders) should be sampled separately in tropical forests, especially for monitoring purposes, and different sets of methods should be combined for each according to their most efficient and best cost-benefit performance.

## Introduction

Arthropods are one of the most abundant groups of organisms in several ecosystems. Therefore, it can be difficult to generate an accurate species list (‘strict inventory’) for a particular area or to estimate patterns of species abundances (‘community characterization’) for comparisons among different assemblages (Longino et al., 2002; Scharff et al., 2003; Barlow et al., 2007; Cabra-García et al., 2012). This task is even more challenging in tropical forests, which account for 17% of the land (Whittaker, 1975) but harbor disproportionately more species than any other terrestrial ecosystem (Gaston, 2000). Therefore, logistic support, financial costs, time and the availability of adequate sampling methods are the main limitations for sampling biodiversity in tropical forests (Gardner et al,. 2008).

Structured inventories incorporate key features of both strict inventories and assemblage characterizations (Oliver & Beattie 1996; Longino & Colwell, 1997) and are highly desirable for revealing the general distribution and the relative abundance of species across different scales and sites. Structured inventories aim to generate accurate species lists of given areas, but also providing reliable estimates of species abundance, which is a key factor for assemblage characterizations (Fisher, 1999; Cardoso, 2009; Tourinho et al., 2014). Structured inventories normally use a combination of several sampling methods and have become an interesting alternative approach for monitoring arthropods or using arthropods as indicators of ecological change and ecosystem dynamics (Souza et al., 2012).

The use of several sampling methods is often required to produce a reliable estimation of species richness and composition (e.g., Coddington et al., 1991; Bonaldo et al., 2009). Although sampling method performance may vary among different taxonomic groups (see Gardner et al., 2008), in tropical forests many different taxonomic groups have been usually sampled together, using a combination of selected sampling methods (e.g., Bonaldo et al., 2009) and requiring massive sampling effort. Given a sufficiently high sample effort, the combination of several sampling methods should adequately represent both species richness and composition (Azevedo et al., 2014). However, the use of several sampling methods is not necessarily optimal for improving monitoring and evaluation programs (Gardner et al., 2008; Porto et al., 2016). Given the costs, monitoring programs often seek an optimum balance between sampling effort and the time consumed to achieve their goals (Souza et al,. 2012). Therefore, the reduction of sampling effort based on the redundancy and complementarity of sampling methods was suggested for invertebrate surveys in several sites in the Amazon basin (Souza et al., 2012; Tourinho et al., 2014; Porto et al., 2016).

We used the opportunity to sample arachnids (mostly spiders and harvestmen) at the Porto Urucu petroleum/natural gas production facility to address the following two questions: (1) Can sampling methods and protocols developed for spiders be reliable for harvestmen? Specifically, we assessed whether the same sampling methods (nocturnal search, litter samples and beating tray) yielded different sets of spiders and harvestmen species and different estimates of local species richness; and (2) What is the most effective single sampling technique for estimating two different taxonomic groups (spider and harvestmen assemblages) in this tropical forest? To examine ecological correlates of spider distribution patterns, we also used a guild classification approach (Dias et al., 2010), which provides a useful framework for describing and analyzing the spider assemblage structure.

## Material and Methods

### Study site

The present study was conducted in Coari municipality, state of Amazonas, Brazil, at Porto Urucu, a petroleum/natural gas production facility belonging to Petrobras S.A. The petroleum facility is located on the right bank of the Urucu River, Solimões River basin, at 4°30’S, 64°30’W and 650 km west of Manaus (Fig. 1). The region is mostly covered by dense upland (‘*terra firme*’) rain forest with uniform canopy, presenting a low diversity of lianas and epiphytes (Lima Filho et al., 2001). About 913 plant species have been recorded and notable changes in the vegetation structure occur in areas with poor soil drainage or in natural or artificial forest gaps (opened for natural gas and oil prospecting and exploiting). Natural gaps are formations produced in the forest matrix from fallen trees or large branches from the canopy, exposing the forest ground to direct solar radiation; whereas artificial gaps were created by the removal of trees and soil materials for the construction or maintenance of the Porto Urucu road network. For this study, records were made only in artificial gaps.

**Fig. 1.**
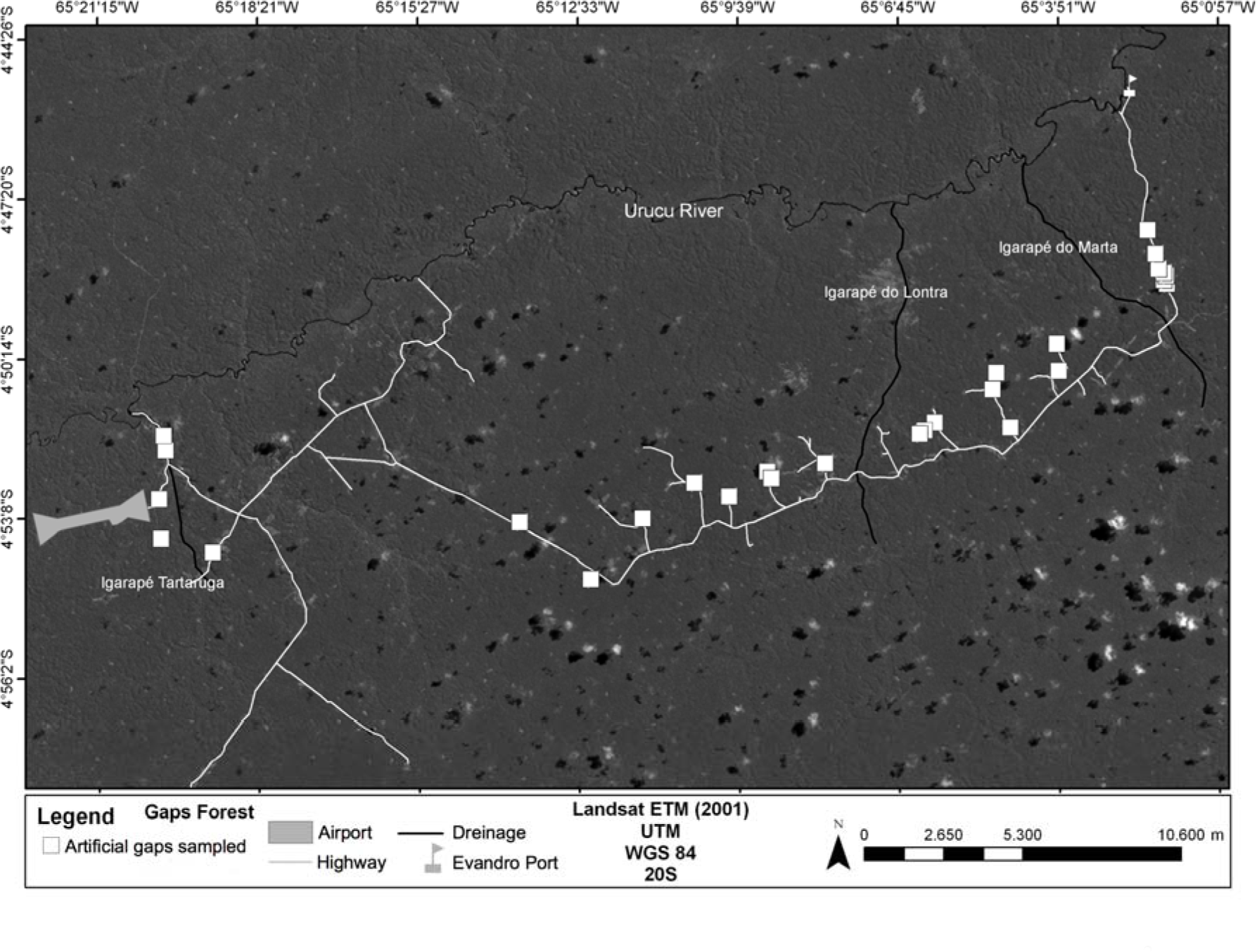
Map of the Porto Urucu sampling site. The squares represent the 33 plots of 300 m^2^ sampled.

### Data collection, species identification and costs

We established our plots in 33 artificial forest gaps. Three collectors sampled the arachnids (see Dias & Bonaldo, 2012, Table 4 for more details) between July and November of 2006. Ten 1 m^2^ samples of litter were sampled by Winkler traps at each one of the collecting plots, quadrats were randomly selected. The invertebrates were extracted from the litter through a 5 mm mesh sieve. The sieved litter was placed in Winkler traps for 48 h. The invertebrates migrate from the suspended litter sample and eventually fall into the pot partially filled with alcohol at the bottom of Winkler extractor. Six to eight beating trays and nocturnal searches were used for sampling in each of the 33 artificial forest gaps. The number of times beating and nocturnal searches were applied have varied from 6-8 between plots, but they never varied among methods into the plot.

The beating tray consisted of a 1 m² white cloth frame placed under the vegetation, which comprised a bush, a shrub or a small leafy branch randomly selected by the collector. The vegetation was vigorously hit with a stick until the invertebrates fall down and the arachnids were hand sorted and stored in a vial containing 80% alcohol. Each sample was a composite of one hour’s sampling (about 20-30 plants sampled).

Each nocturnal search sample was composed by 30 m transect solo walked slowly, searching with a headlamp and hand-searching for the arachnids (looking up and looking down) for 1 hour. Therefore, beating trays and nocturnal searches were standardized by time (1 h), within an area of 300 m^2^ to control sampling effort (see Davies, 1986; Coddington et al., 1991; Pinto-da-Rocha & Bonaldo, 2006 for the methods’ definition and description). The differences between the number of beating and nocturnal search per plot did not bias our further comparisons because sampling effort was always similar within plots. Samples of the same method were summed resulting in one sample for each sampling method per plot. Voucher specimens of both orders are deposited in the arachnological collection of the Museum Paraense Emílio Goeldi - MPEG (A.B. Bonaldo, curator), Belém, Pará, Brazil.

Each individual sampled was carefully examined by the experts in taxonomy and systematics of spiders and harvestmen (Ana Lúcia Tourinho, Ricardo Pinto-da-Rocha, Alexandre Bonaldo and staff, Antonio Brescovit and staff, Erika Buckup and staff). They were dissected and studied using both stereomicroscope and light microscope, and we used the somatic characters and the sexual characters to identify our material. Whenever possible identifications to species level were provided; otherwise morphospecies were defined. Only adult individuals were used because most juveniles cannot be adequately identified since their sexual characters are not fully developed, which is extremely important for accurate identification of both harvestmen and spiders. Those taxa in poor taxonomic state of knowledge were not included in any specific genera to avoid misinterpretation (e.g. Cosmetidae sp.1 and Cosmetidae sp.2). For this set of species it is only possible to confirm whether the species belong to the same genus or not after a taxonomic revision. All the potential new genera and species, very common in any inventory undertaken in the Amazon, were referred to as genus or species followed by their number (e.g. Gagrellinae, genus 1 sp.1).

We followed Souza et al. (2012), Gardner et al. (2008) and Porto et al. (2016) to evaluate the project costs. We considered financial costs and time spent collecting in the field, during sorting and identifying in the lab. The summed time of all sampling techniques together was set up as maximum effort (100%) and we calculated the fractions of maximum effort for the three methods employed for each combination of sampling techniques.

### Data analyses

For both spiders and harvestmen, we estimated the species richness of the site based on data from each sampling method, equally sampled among the 33 plots. We also quantitatively compared the relative sampling efficiencies of the nocturnal search, litter sample and beating tray approaches, and the assemblage species compositions between sampling methods for each arachnid order separately. The raw data is available in the supplementary material (Table 1, 2 and 3).

**Table 1.** Optimal combinations of samples per method for each site given 50%, 80% or 100% of species sampled at Porto Urucu petroleum facility, Amazonas, Brazil.

|  | Harvestman |  |  | Spiders |  |  |
| --- | --- | --- | --- | --- | --- | --- |
| method | 50% | 80% | 100% | 50% | 80% | 100% |
| Beating | 1 | 1 | 25 | 12-13 | 33 | 33 |
| Nocturnal | 2-3 | 11-13 | 33 | 5 | 11-12 | 33 |
| Winkler | 0 | 1 | 5 | 0 | 2 | 33 |

**Table 2.**
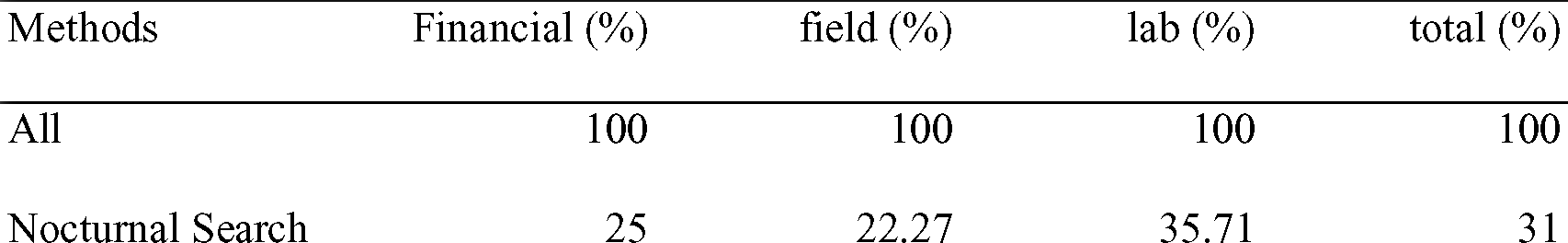
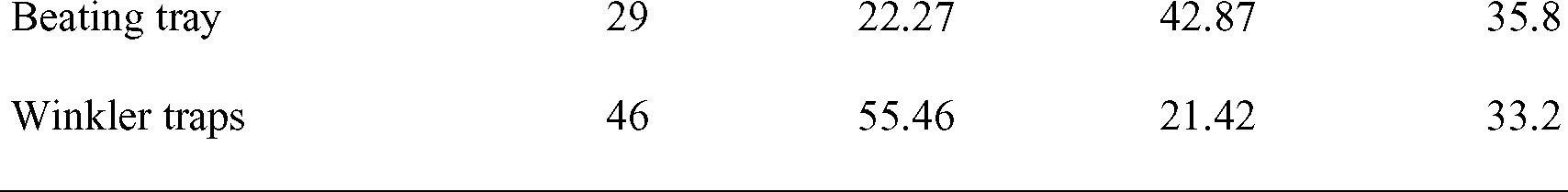
Financial and time costs spent in the field and lab for each method.

We used the Chao1 index to estimate the species richness of the 33 plots for both spiders and harvestmen (Chao 1984). The 95% confidence intervals (CIs) associated with these estimates of species richness were also calculated based on 200 bootstraps (Colwell, 2012). For this analysis, we pooled the data for all the replicate traps regularly distributed among the 33 sampling plots within each sampling method. The Chao1 richness estimator provides a conservative minimum estimate of the number of species that are present, accounting for the non-collected species in the samples (Colwell & Coddington 1994).

Comparative analyses of sampling method performance can be biased by variations in sampling intensity between techniques. Even with standardized sampling, biodiversity measures remain sensitive to the number of individuals and the number of samples collected (Gotelli & Colwell 2001). Rarefaction methods calculate the expected number of species based on a random subsample of the data, making the comparisons among sampling methods more reliable. However, the sampling methods differed greatly in the number of individuals they accumulated. Therefore, we used sample-based and incidence-based (the average species accumulation curves based on the abundance of individuals) rarefactions to compare the number of species between sampling methods (Gotelli & Colwell, 2001). We used analytical methods to generate valid CIs for the rarefaction curves, which do not converge to zero at the maximum sample size (Colwell et al., 2004). Calculations and simulations were performed using EstimateS (version 8.2) (Colwell, 2012). We also estimated whether the species accumulation curve reached the asymptote, by calculating the first derivative at the last sample using the package BAT in the R environment (Cardoso et al., 2015).

To evaluate the composition similarity between sampling methods, we calculated the Jaccard similarity index between each pair of collection methods, accounting for the bias toward small values, as this index does not take into account (rare) shared species that were not represented in either of the two sample collections (Chao et al., 2005). Therefore, the modified Jaccard index is not upper bounded, and we used 1,000 random bootstrap samples to calculate 95% CIs for this index. When the CIs encompass 1.0, we can accept the null hypothesis that the two sampling methods share a similar group of species. Calculation of the adjusted Jaccard similarity and construction of the bootstrapped CIs were done using EstimateS (version 8.2) (Colwell 2012). We also compare compositional similarity between sampling methods using the Bray-Curtis index to account for differences among species relative frequencies.

We used the framework proposed by Cardoso (2009) to optimize the number of samples per method. This analysis permutes the data matrix focusing on a combination of sampling methods that maximizes a target number of species. We showed the better combination of methods and number of samples per method to achieve 50%, 80% and 100% of species sampled at the Urucu site. These analyses were calculated using the function ‘*optim.alpha*’ of the BAT package (Cardoso et al. 2015), and were based on 1000 permutations.

To show the composition and identity of species sampled by each method we created indirect ordination graphs, where assemblage composition of the plots per method were ordered based on the Bray-Curtis dissimilarity index. Given the high number of spider species, we showed only the ordination figures for spider families.

Spiders are incredibly diverse, however, and the level of classification detail depends on the information needed in each study. Therefore, guild classification is an interesting approach to exploring the general patterns among the sampling techniques as different families within same guild can present similar roles in the ecosystems (Cardoso et al., 2011). This guild classification is based on natural history information obtained through direct observation of individuals hunting, resting, building webs, carrying egg-sacs, running, stalking and ambushing (Dias et al., 2010). This classification is not exclusively based on family level; it was revised to reflect the natural history below family level. An inferential test to assess possible differences in guild abundances among sampling methods was made by a permutational multivariate analysis of variance (PERMANOVA, Anderson, 2001), using the function ‘*adonis*’ in the package ‘Vegan’. For this analysis we did not used the Jaccard similarity index accounting for the bias toward small values, as the sampling representation at guild level was reasonably high. The PERMANOVA was based on the Bray-Curtis distance measurement. This analysis was performed in R (version 2.14) (R Development Core Team 2011) using the ‘Vegan’ package (Oksanen 2012).

## Results

We sampled 2,139 harvestmen and 3,786 spiders distributed into 26 harvestmen species and 625 spider species, respectively. The beating-tray method sampled 1,236 harvestmen (7 families, 13 species) and 1,969 spiders (28 families, 417 species). The nocturnal search method sampled 667 harvestmen (9 families, 24 species) and 1,537 spiders (30 families, 357 species). The Winkler traps sampled 236 harvestmen (9 families, 13 species) and 280 spiders (15 families, 91 species). Given the different sampling methods used, the species sampled included arboreal, soil and litter specialist species (supplementary material).

For harvestmen, the nocturnal search method sampled significantly more species than either the Winkler or beating tray methods, as can be seen by examination of the 95 % confidence intervals (Fig. 2). The nocturnal search was also more effective, sampling more species and fewer individuals (Fig. 3). Relatively fewer harvestmen individuals were sampled using the Winkler method (Fig. 3). However, the Winkler trap method seems to be more efficient than the beating tray, as the number of species sampled by each method is similar given a similar number of samples (Fig. 2 and 3). The sampling effort was sufficient to sample a reasonable proportion of the harvestman richness at the Urucu site. Beating tray, nocturnal search and Winkler sampled 87%, 91% and 43%, respectively, of the total number of species expected to be sampled with each method based on Chao1. Despite the lower proportion sampled with Winkler, the slopes at the end of the accumulation curves for all methods (beating, nocturnal search and Winkler) were close to zero (< 0.009 in all cases).

**Fig. 2.**
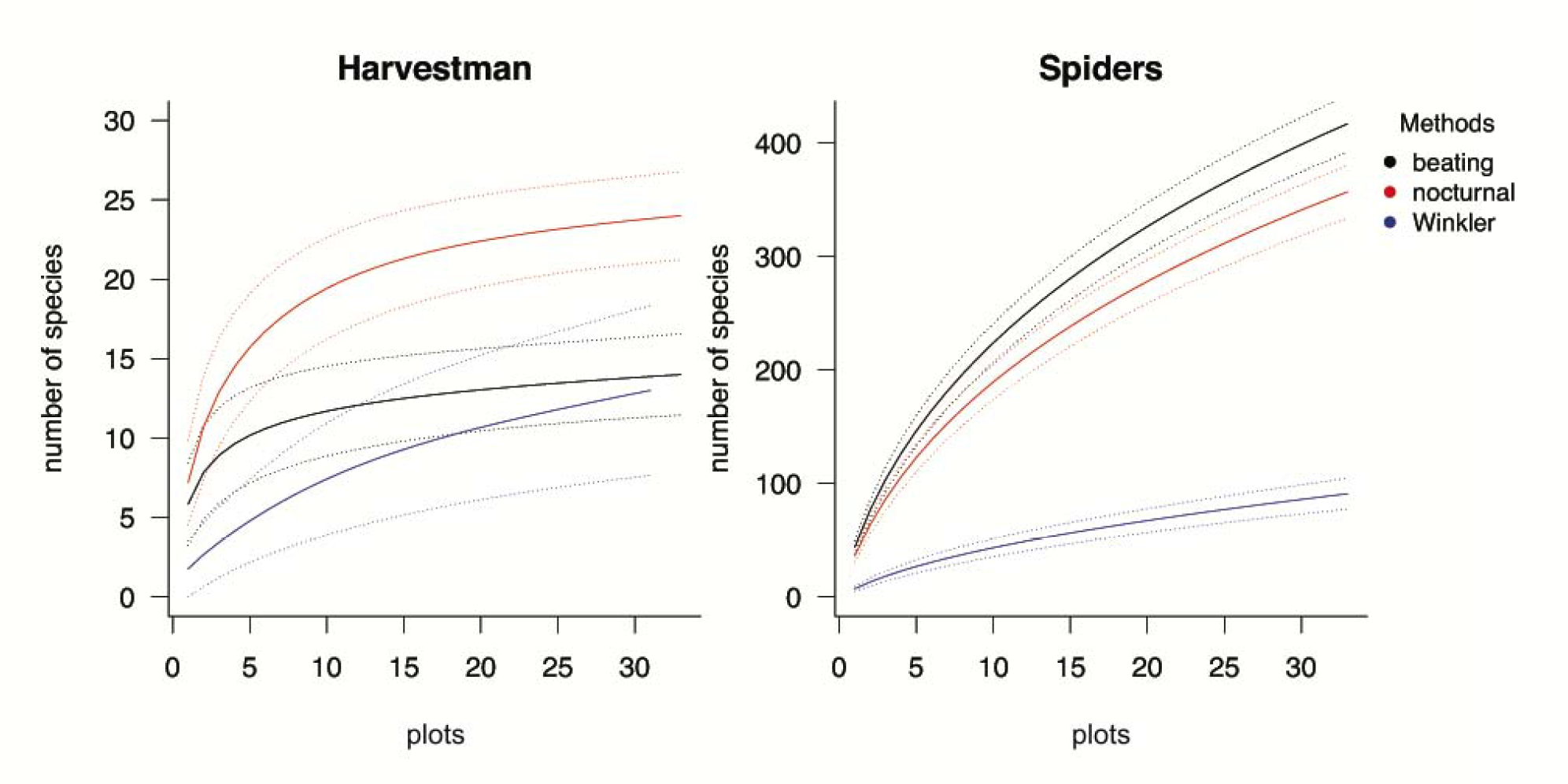
Sampled based rarefaction curves for harvestmen and spiders. Fitted dotted lines indicate 95% confidence intervals; tracing dotted lines indicate number of species estimated.

For spiders, the Winkler method sampled far fewer species than either the beating tray or nocturnal search (Figs. 2 and 3). In absolute terms, the beating tray method collected more species than the nocturnal search per plot (Fig. 2). However, the rarefaction curves of the two methods are not different based on the number of individuals sampled (Fig. 3). Unlike the situation for harvestmen, none of the species accumulation curves for spiders were close to being saturated (Figs. 2 and 3). The slopes at the end of the accumulation curves for all methods (beating, nocturnal search and Winkler) was ~0.1 in all cases, meaning that if a new sampling exercise was carried out we expect to find ~3 new species using the beating tray, ~2.5 species during nocturnal search and ~0.6 species in a new Winkler sample. Beating tray and nocturnal search sampled ~60% and Winkler sampled ~43% of the total number of spiders species expected to be sampled by each method based on Chao1.

**Fig. 3.**
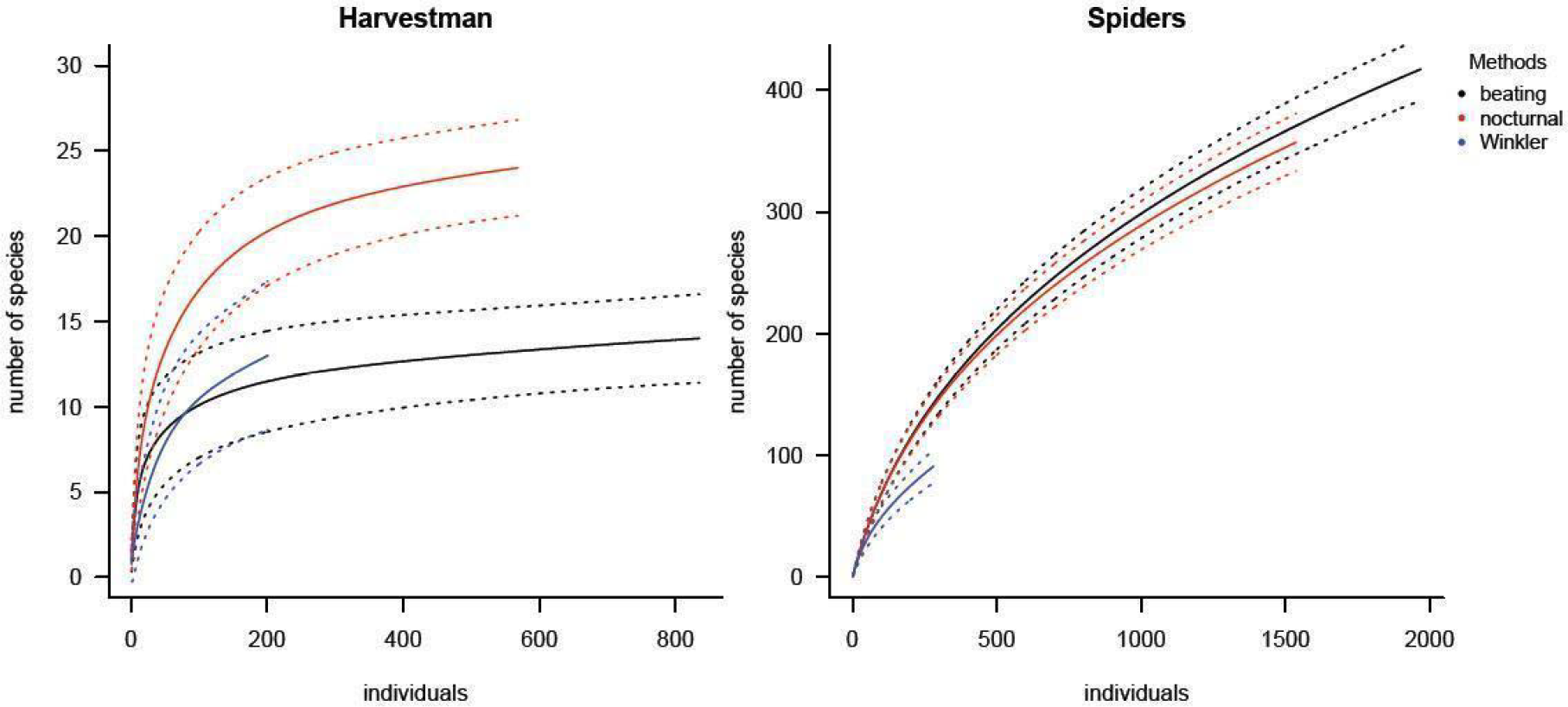
Sample based rarefaction curves for harvestmen and spiders. Fitted dotted lines indicate 95% confidence intervals; tracing dotted lines indicate number of species estimated.

**Fig. 4.**
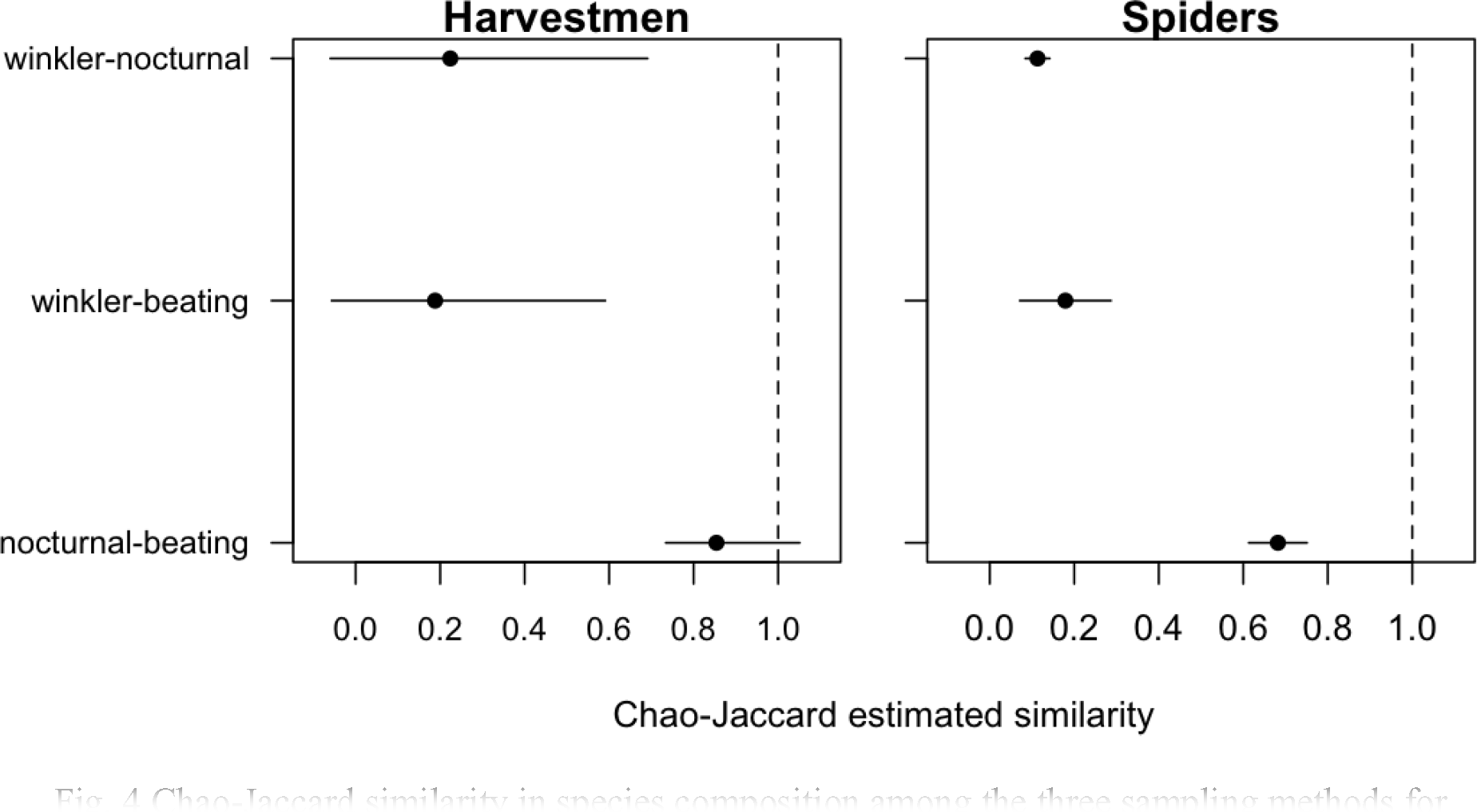
Chao-Jaccard similarity in species composition among the three sampling methods for harvestmen and spiders, adjusted for unsampled species. The point represents the mean similarity and the line around the point the 95% confidence intervals.

The three techniques generally sampled different sets of species at the plot scale for both harvestmen and spiders (Fig. 4). The only exception was for the harvestman assemblage sampled with nocturnal search and beating tray. In this pairwise comparison, the adjusted compositional similarity was around 85% with the 95% CI including 1 (Fig. 4). The spider assemblage compositions sampled by nocturnal search and beating tray were also more similar, although the 95% CI was less variable. The other pairwise comparisons for both harvestmen and spiders assemblages composition have far fewer species in common. While each sampling method samples individuals at different ratios, the same general pattern was retrieved using relative abundance data (Bray-Curtis index). Both harvestman and spider assemblages sampled with nocturnal search and beating tray were more similar (~36% for harvestmen and ~44% for spiders) than other sampling methods comparisons (7-11% for harvestman and 3-6% for spiders).

At the site scale, the harvestman species compositions sampled by nocturnal search, beating tray and Winkler were redundant (Fig. 5), but the nocturnal search sampled more exclusively harvestman species than any other method. While the nocturnal search collected seven exclusive species, the Winkler method sampled only one exclusive species and all species sampled with the beating tray were also sampled by the other methods. This result is also expressed in the optimization of sampling methods, while a combination of 11 or 12 nocturnal search coupled with one beating tray and one Winkler sample may account for 80% of harvestman species sampled (Table 1).

**Fig. 5.**
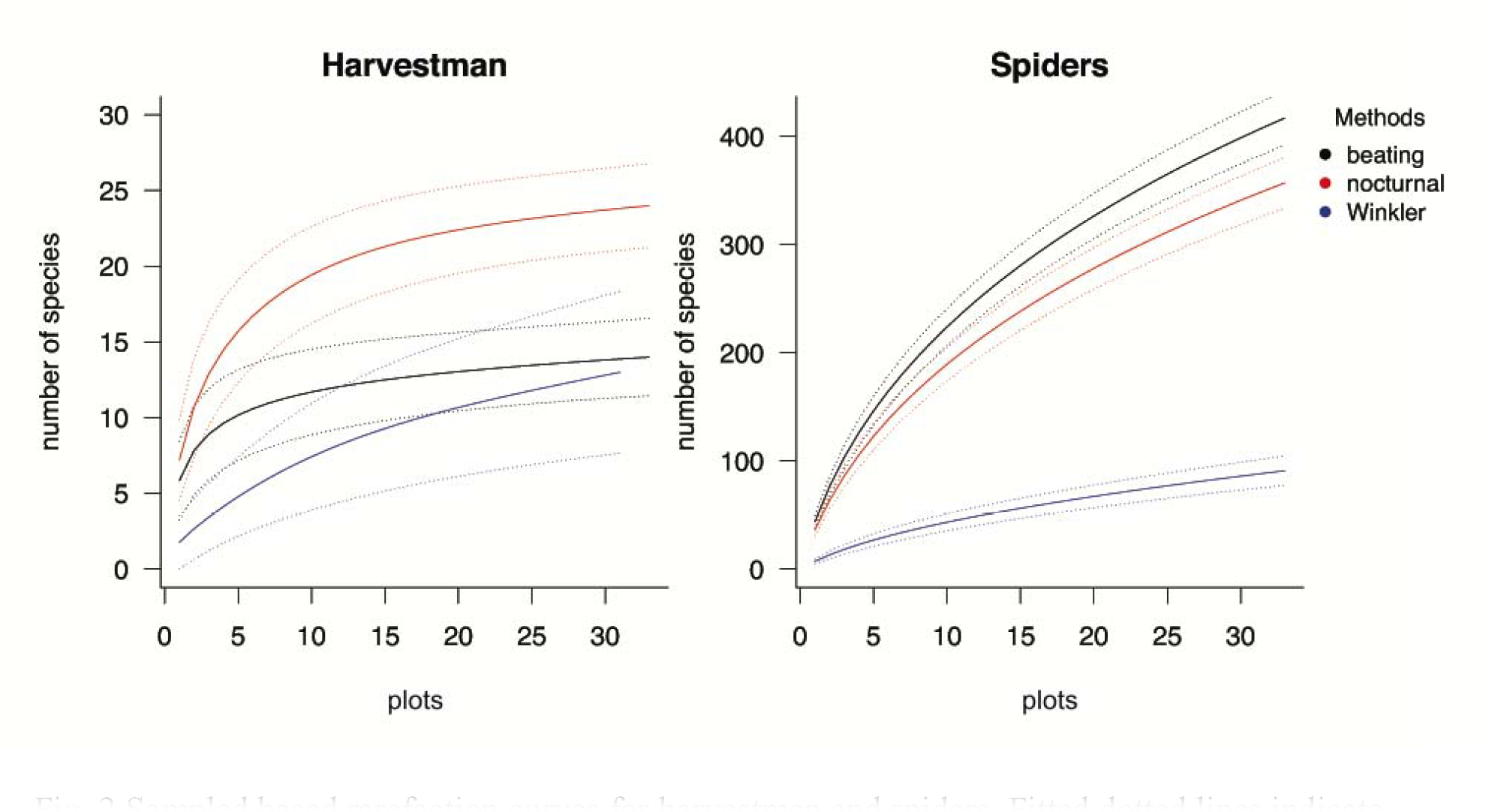
ChaDistribution of harvestman species sampled by each technique among the study plots at Porto Urucu, Amazonas, Brazil. The colors scheme is the same as in Fig. 2 (black bars represent the species sampled with beating tray, red bars the species sampled during nocturnal search and blue bars the species sampled with Winkler traps). Each column represents one sampling plot. The bar size represents the relative abundance standardized by the maximum number of individuals recorded in one plot.

The spider assemblage at site scale showed a markedly different picture. Each sampling method collected a set of exclusive species (Fig. 6). Figure 6 shows the spider families collected by each method. Given the large number of species sampled it is infeasible to show all of them on a single plot. The sampling optimization procedure suggests that Winkler samples are more redundant to sample 50% of harvestman species, but this is happening because Winkler samples accumulates species at lower rates compared with other methods (it is less efficient). To detect 80% of spiders species (498 species), all beating samples, almost half of nocturnal search, and two Winkler samples is necessary (Table 1). The distribution pattern of spider guilds among sampling methods was similar to the distribution of species or families (Fig. 7). Each sampling method favored specific set of spider guild compositions (PERMANOVA, r^2^ = 0.52, *F* = 53.235, p < 0.001), although the differences are more related with guild relative abundance than occurrence (Fig. 7).

**Fig. 6.**
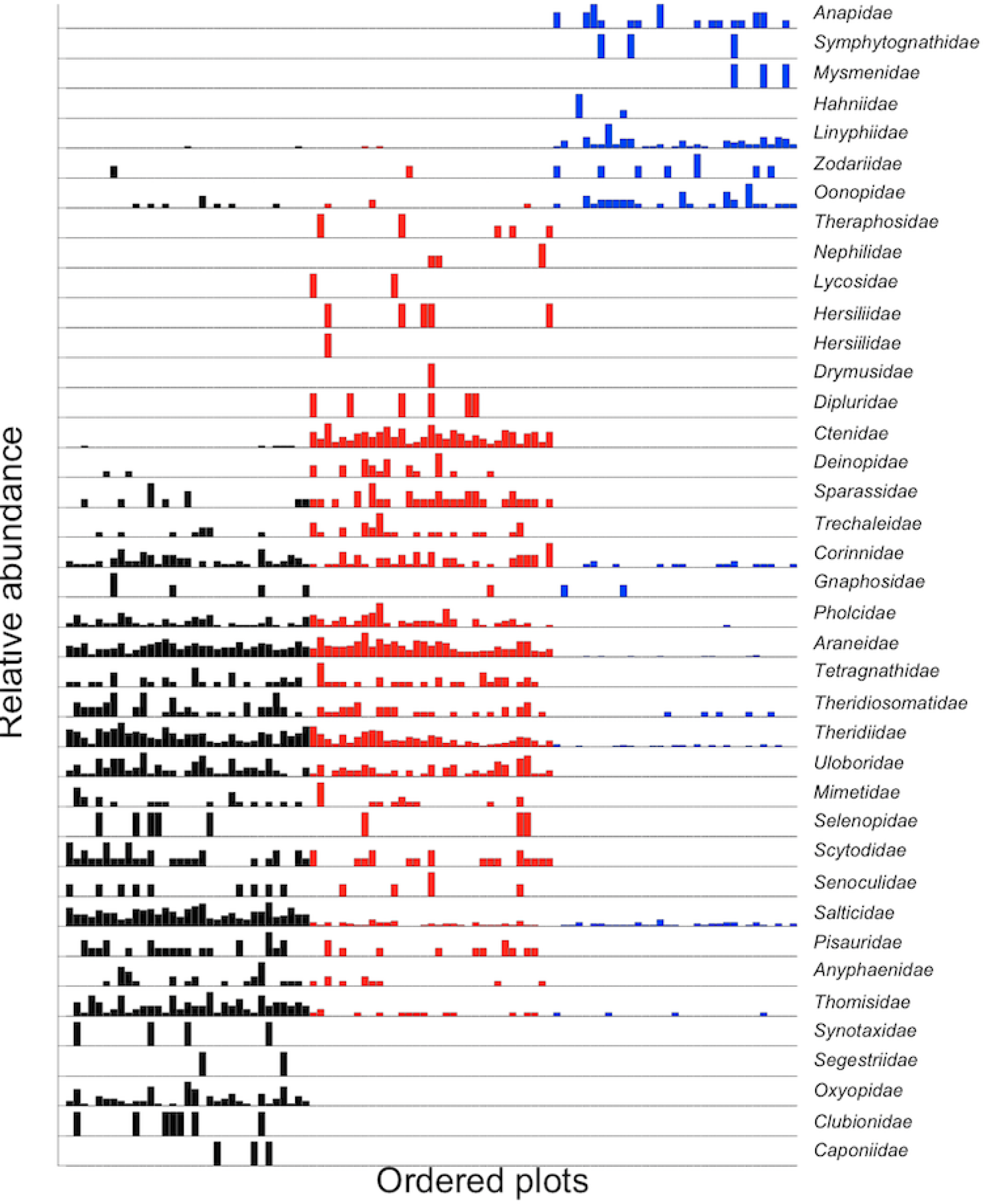
ChaDistribution of spiders families sampled by each technique among the study plots at Porto Urucu, Amazonas, Brazil. The color scheme is the same as in Fig. 5 (black bars represent the species sampled with beating tray, red bars the species sampled during nocturnal search and blue bars the species sampled with Winkler traps). Each column represent one sampling plot, and the bar size represents the relative abundance standardized by the maximum number of individuals recorded in one plot.

**Fig. 7.**
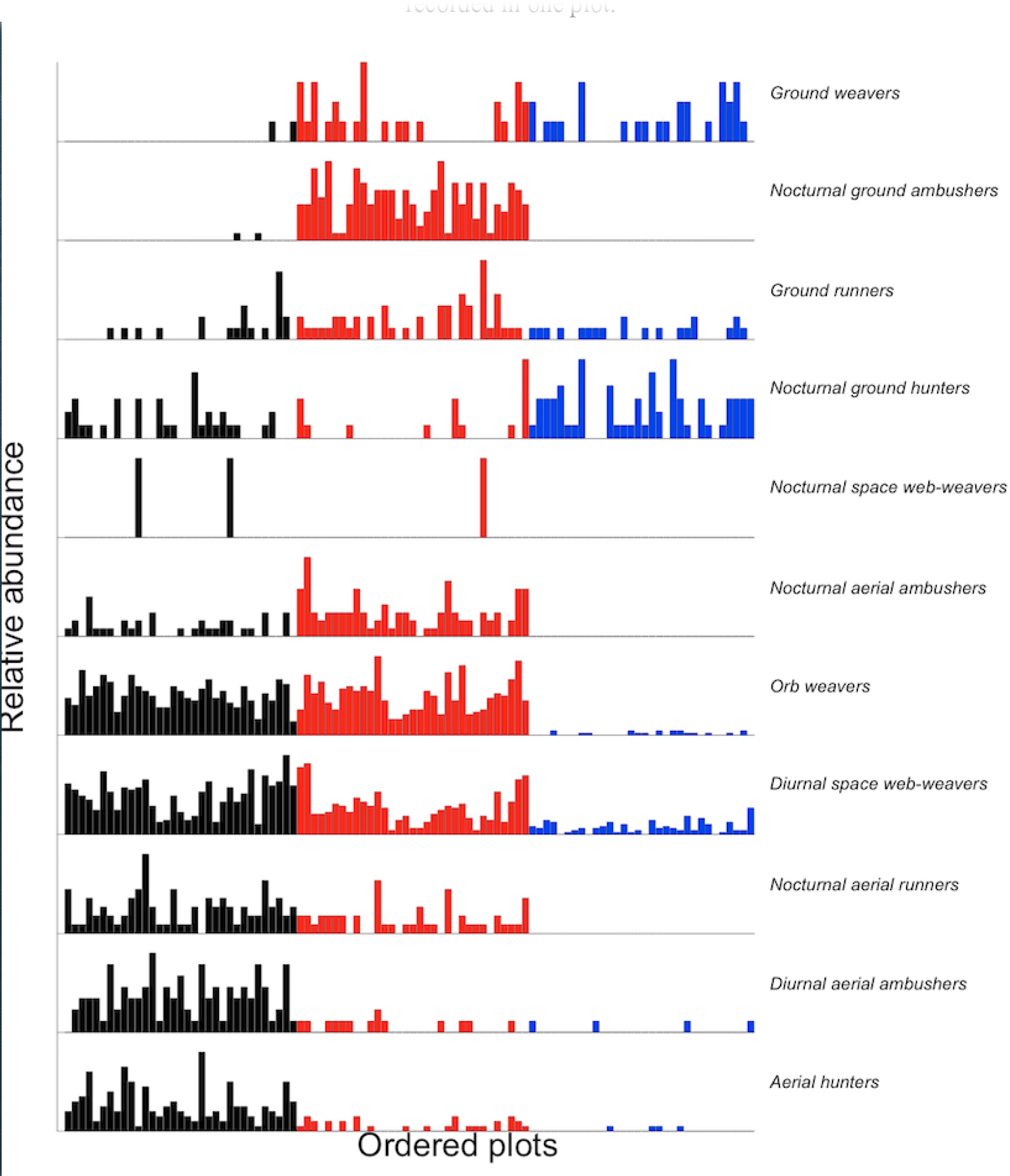
ChaDistribution of spider guilds sampled by each technique among the study plots at Porto Urucu, Amazonas, Brazil. The color scheme is the same as in Fig. 5 (black bars represent the species sampled with beating tray, red bars the species sampled during nocturnal search and blue bars the species sampled with Winkler traps). Each column represent one sampling plot, and the bar size represents the relative abundance standardized by the maximum number of individuals recorded in one plot.

Winkler traps consumed nearly half of the financial costs and took most of the time taken in the field (Table 2). This method consumed 55.46 % of time for the study dedicated for the survey in the field. Nocturnal search is the cheapest and faster method, only 25% of financial resources were consumed and 31% of the total time was dedicated for the study. Beating consumed only 29% of the financial resources; however it was as fast as nocturnal sampling in the field, but consumed 42.87 % of time in the lab sorting and identifying the material surveyed.

## Discussion

The efficiency of the methods used to simultaneously sample different taxonomic groups of arachnids have not been explored or compared before. In our study the most striking difference between spider and harvestmen inventory data is that none of sampling methods got close to achieving the asymptote of accumulation curves for spiders. However the same sampling effort was sufficient to describe the species richness of harvestmen at the Urucu site. Therefore, a reasonable inventory of harvestmen is then accomplished with less effort and fewer methods than would be needed for recovering similar results with spiders. Based on Chao1 estimates 89% of harvestman species were sampled, while only 66% of spider species were detected with all methods combined. The higher richness, abundance, complexity and diversity of the spider guild accounts for the need for several sampling methods and several replicates that are not necessary for a reasonable inventory of harvestmen in tropical forests.

The nocturnal search method was the most efficient method for accessing the species richness of harvestmen in the study area, while Winkler traps and beating were less efficient and were redundant with the nocturnal search method. These results are corroborated by two previous studies comparing the efficiency of sampling methods for this group in three different areas in the Amazon basin (Tourinho et al., 2014; Porto et al., 2016). Differently, the beating tray method collected more spider species and more individuals than either the nocturnal search or Winkler methods. In that regard, our findings differ from other studies (Coddington et al., 1991; Cardoso, 2008; Azevedo et al., 2014) and cast doubt on the claim that nocturnal search is the most productive method for accessing spider richness and should be used alone in tropical forests (Azevedo et al., 2014). Based on our results, the performances of the beating tray and nocturnal search approaches are equivalent for describing the local richness of spiders in the Urucu basin and both methods should be used simultaneously to describe it in the Amazon basin.

While for harvestmen the species composition for the area using the maximum effort (three methods combined) can be virtually summarized solely by nocturnal search, for spiders each sampling method sampled a different set of species. Both the nocturnal search and beating tray approaches sampled almost the same number of species of spiders; however, each method recorded a different set of species and at different relative abundance. Similar richness and different species composition between nocturnal search and beating tray suggests that these two methods have little redundancy in this tropical forest and richness estimates based on only one of these sampling methods will result in very biased results. Whereas in some cases two methods sampled virtually the same set of species (i.e. only one harvestman species was exclusively sampled by Winkler or beating tray), the compositional similarity between sampling methods using relative abundances of species (Bray-Curtis index) were relatively lower for all methods comparisons. These results suggest that each method sample different number of individuals, despite species identity. Despite the differences in species composition and abundance found at the plot level, the same pattern can be detected at the site scale. For harvestman, the redundancy of sampling methods is very high, with only one species exclusively sampled by Winkler or beating tray, while the majority of the species was sampled by nocturnal search alone. These results suggest that complementarity among sampling methods can be detected in small-scale inventories for spiders, while redundancy between sampling methods is strong for harvestman inventories.

The pattern of spider guilds composition among sampling methods was as expected for a typical Neotropical forest. Winkler traps recorded mainly species with small body lengths that live on the ground or in the leaf litter, such as Ground weavers, Ground hunters and Nocturnal Ground hunters, which are among the spiders that are normally accessed by this technique (e.g., Anapidae, Hahniidae, Oonopidae, Tetrablemmidae, Zodariidae and Zoridae). Also observed in this habitat are the large ground spiders such as Ctenidae and several Mygalomorphae that are too big to pass through the sieve’s grid. These spiders are much more likely to be sampled by nocturnal search.

The higher spider abundance recorded during the nocturnal search was due to the inclusion of nocturnal ground ambushers, nocturnal aerial ambushers, orb weavers and diurnal space web weaver species. Despite being diurnal, webs produced by these spiders remain intact or in pieces during the night. Given that spiders are often found in retreats near the webs and since the webs are very conspicuous at night during light focal search, this guild can be well documented with nocturnal sampling. Beating was usually sampled in the morning, and the presence of nocturnal species can be explained by the fall of spiders that were resting or hidden in the bushes.

Nocturnal search was the most effective method to sample harvestmen, and both nocturnal search and beating tray were the most productive sampling methods for spiders. While both methods provide a comprehensive coverage of species spider species that are present at a site, they may not be the best option for sampling abundances or biomass per unit of area for most species. Nocturnal search and beating vegetation can be standardized by area, but given that Winkler virtually remove all the microhabitat available (litter), it may offer more realistic informations on species abundances and biomass per unit of area. However, this feature must be interpreted with caution. Given that leaflitter samples are passed through a sieve before it is hanged to dry out, and it also has a second sieve inside the Winkler, this method is selecting species by size in advance, thus it may be efficient to cover biomass and abundance for individuals which are often small-sized species with cryptic behavior (Krell et al. 2005).

Harvestmen species possess low dispersal capability and are generally highly endemic (Pinto-da-Rocha et al., 2007), while spider species tend to disperse to greater distances, even dispersing across continents through ballooning (Foelix, 2010). Despite the fact that several spider species also have low dispersal capability, notably some clades are typically encountered in leaf litter (e.g., Oonopidae), and thus might be expected to present micro-distributed patterns even in Amazonia. The differences in species dispersal abilities between harvestmen and spiders described above often result from different species distributions in the forest. Therefore, the methods and protocols developed for spiders are not always similarly successful for harvestmen.

Our results were different in terms of efficiency and congruence for each of the groups sampled. The financial and temporal investment for collecting and sorting spiders is more than twice the investment needed for creating an inventory for harvestmen. Beating tray is the second most expensive method, it consumed the same time as nocturnal search in the field and 42% of the time processing the material in the lab. However, as there is a high complementarity between spider species sampled with nocturnal search and beating tray, these methods should be used together if a more detailed picture of spider diversity is needed. To get nearly 80% of spider species sampled all beating trays and about half of nocturnal search needed to be employed.

Multiple methods and several replicates are used to sample spiders in many studies and the accumulation curves are also still not asymptotic (Coddington et al., 1991; Bonaldo et al., 2009; Cabra-García et al., 2010; Azevedo et al., 2014). Therefore, a traditional spider sampling protocol is usually not the most efficient strategy for studies of assemblage associations with environmental variables and ecological impact. No single sampling method can adequately sample a full representation spider assemblages. However, depending on the question posed, an especific combination of sampling methods may be more adequate than others. Our results suggest that a combination of Winkler and beating vegetation may give a more comprehensive information about abundance and species richness, while Winkler plus nocturnal search will give a good estimative of whole species assemblage, whilst reducing costs. Further tests are needed to compare abundances and biomass efficiency among methods. It is nearly impossible to assess a complete spider species inventory in the tropics, so larger protocols demand higher financial costs, increasing time spent in both the field collecting and in the lab processing. An interesting option to overcome this problem is using spider guilds in long term monitoring studies. For the Urucu forest, for example, an efficient way to monitor a large number of spider’s species would be using only beating tray, but Ground weavers and Nocturnal ground ambushes will be underestimated. Conversely, beating and Winkler traps are redundant, temporal and financially costly for harvestman assemblages, and nocturnal sampling provides a reasonable view of the harvestman assemblage at lower cost.

It is clear that a protocol developed for spider surveys is inadequate for harvestmen in terms of efficiency, congruence and financial cost, especially for monitoring or ecological impact studies. We propose that different taxonomic groups such as harvestmen and spiders should not be surveyed simultaneously in Tropical forests. Nocturnal search is still the most efficient method to sample harvestmen at lower cost and for optimum results. We still need future studies to test and elaborate an optimum protocol for spiders in the Amazon tropical forests. In this paper however, we present an economic and efficient alternative protocol, composed of active nocturnal search and beating tray, which saves almost 50% of the costs.

## Acknowledgments

This work was supported by Conselho Nacional de Pesquisa e Desenvolvimento Tecnológico – CNPq (NFLM #130775/2011-8), (ABB PQ#304965/2012-0), Rede CTPetro Amazônia – Fundo Setorial do Petróleo (FINEP/CNPq/PETROBRAS), PPBio-Amazônia Oriental. ALT and FBB are funded by CENBAM/PPBio and RPR is co-funded by FAPESP (BIOTA, 2013/50297-0), NSF (DOB 1343578) and NASA.

## Contribution of Authors

ALT and FBB performed the analysis, generated the figures, tables and wrote the paper. ABB, SCD and NFLMH collected the data. ALT, ABB, RPR, NFLMH and SCD identified specimens. SCD generated the map. All authors revised and commented on the text.

